# Recent demography drives changes in linked selection across the maize genome

**DOI:** 10.1101/031666

**Authors:** Timothy M. Beissinger, Li Wang, Kate Crosby, Arun Durvasula, Matthew B. Hufford, Jeffrey Ross-Ibarra

## Abstract

Genetic diversity is shaped by the interaction of drift and selection, but the details of this interaction are not well understood. The impact of genetic drift in a population is largely determined by its demographic history, typically summarized by its long-term effective population size (*N*_*e*_). Rapidly changing population demographics complicate this relationship, however. To better understand how changing demography impacts selection, we used whole-genome sequencing data to investigate patterns of linked selection in domesticated and wild maize (teosinte). We produce the first whole-genome estimate of the demography of maize domestication, showing that maize was reduced to approximately 5% the population size of teosinte before it experienced rapid expansion post-domestication to population sizes much larger than its ancestor. Evaluation of patterns of nucleotide diversity in and near genes shows little evidence of selection on beneficial amino acid substitutions, and that the domestication bottleneck led to a decline in the efficiency of purifying selection in maize. Young alleles, however, show evidence of much stronger purifying selection in maize, reflecting the much larger effective size of present day populations. Our results demonstrate that recent demographic change — a hallmark of many species including both humans and crops — can have immediate and wide-ranging impacts on diversity that conflict with would-be expectations based on *N*_*e*_ alone.

---

The genetic diversity of populations is determined by a constant interplay between genetic drift and natural selection. Drift is a consequence of a finite population size and the random sampling of gametes each generation^1^. In contrast to the stochastic effects of drift, selection systematically alters allele frequencies by favoring particular alleles at the expense of others as a result of their effects on fitness. Researchers often study drift by excluding potentially selected sites^2,3,4^, or selection by focusing on site-specific patterns under the assumption that genome-wide diversity reflects primarily the action of drift^5^.

Drift and selection do not operate independently to determine genetic variability, however, in large part because linkage allows the effects of selection to be wide-ranging^6,7,8^. Linked selection, which refers to the effects of selection at one site on diversity at linked sites^8^, can take the form of hitch-hiking, when the frequency of a neutral allele changes as a result of positive selection at a physically linked site^6^, or background selection, where diversity is reduced at loci linked to a site undergoing selection against deleterious alleles^9^. Recent work in *Drosophila*, for example, has shown that virtually the entire genome is impacted by the combined effects of these processes^10,11,12^.

The impact of linked selection, in turn, is heavily influenced by the effective population size (*N*_*e*_), as the efficiency of natural selection is proportional to the product *N*_*e*_*s*, where *s* is the strength of selection on a variant^8,13,14,15^. The effective size of a population is not static, and nearly all species, including flies^16^, humans^17^, domesticates^18,19^, and non-model species^20^ have experienced recent or ancient changes in *N*_*e*_. Although much is known about how the long-term average *N*_*e*_ affects linked selection^13^, relatively little is understood about the immediate effects of more recent changes in *N*_*e*_ on patterns of linked selection.

Because of its relatively simple demographic history and well-developed genomic resources, maize (*Zea mays*) represents an excellent organism to study these effects. Archaeological and genetic studies have established that maize domestication began in Central Mexico at least 9,000 years bp^21,22^, and involved a population bottleneck followed by recent expansion^23,24,25^. Because of this simple but dynamic demographic history, domesticated maize and its wild ancestor teosinte can be used to understand the effects of changing *N*_*e*_ on linked selection. In this study, we leverage the maize-teosinte system to study these effects by first estimating the parameters of the maize domestication bottleneck using whole-genome resequencing data and then investigating the relative importance of different forms of linked selection on diversity in the ancient and more recent past. We show that, while patterns of overall nucleotide diversity reflect long-term differences in Ne, recent growth following domestication qualitatively changes these effects, thereby illustrating the importance of a comprehensive understanding of demography when considering the effects of selection genome-wide.

## RESULTS

### Patterns of diversity differ between genic and intergenic regions of the genome

To investigate how demography and linked selection have shaped patterns of diversity in maize and teosinte, we analyzed data from 23 maize and 13 teosinte genomes from the maize HapMap 2 and HapMap 3 projects^26,27^. As a preliminary step, we evaluated levels of diversity inside and outside of genes across the genome. We find broad differences in genic and intergenic diversity consistent with earlier results^28^(Figure 1). In maize, mean pairwise diversity (π) within genes was significantly lower than at sites at least 5 kb away from genes (0.00668 vs 0.00691, *p* < 2 × 10^−44^). Diversity differences in teosinte are even more pronounced (0.0088 vs. 0.0115, *p* ≈ 0). Differences were also apparent in the site frequency spectrum, with mean Tajima’s D positive in genic regions in both maize (0.4) and teosinte (0.013) but negative outside of genes (−0.087 in maize and −0.25 in teosinte, *p* ≈ 0 for both comparisons). These observations suggest that diversity in genes is not evolving neutrally, but instead is reduced by the impacts of selection on linked sites.

**Figure 1:**
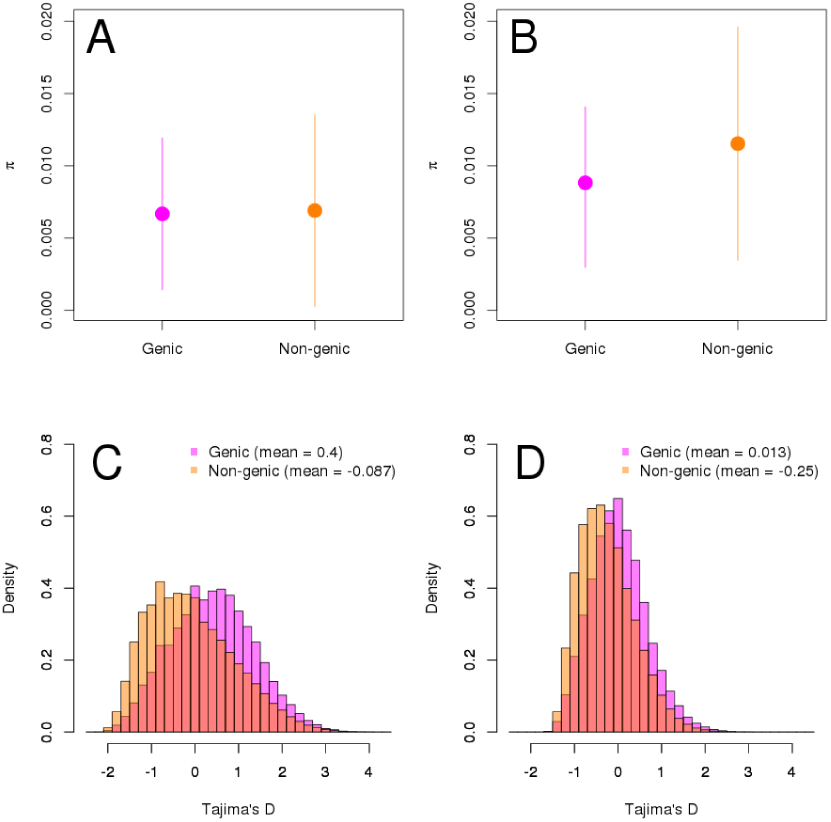
**A** and **B** show mean pairwise diversity *π*, ± one standard deviation, while **C** and **D** depict and Tajima’s D in 1kb windows from genic and nongenic regions of maize and teosinte.

### Demography of maize domestication

We next estimated a demographic model of maize domestication (Figure 2). To minimize the impact of selection on our estimates^29^, we only included sites >5kb from genes. The most likely model estimates an ancestral population mutation rate of *θ* = 0.0147 per bp, which translates to an effective population size of *N*_*a*_ ≈ 123, 000 teosinte individuals. We estimate that maize split from teosinte ≈ 15, 000 generations in the past, with an initial size of only ≈ 5% of the ancestral *N*_*a*_. After its split from teosinte, our model posits exponential population growth in maize, estimating a final modern effective population size of *N*_*m*_ ≈ 370, 000. Although our model provides only a rough approximation of migration rates, we included migration parameters during demographic inference because omitting these could bias our population size estimates. We observe that maize and teosinte have continued to exchange migrants after the population split, with gene flow from teosinte to maize was *M*_*tm*_ = 1.1 × 10^−5^ × *N*_*a*_ migrants per generation, and from maize to teosinte we estimate *M*_*mt*_ = 1.4 × 10^−5^ × *N*_*a*_ migrants per generation.

**Figure 2:**
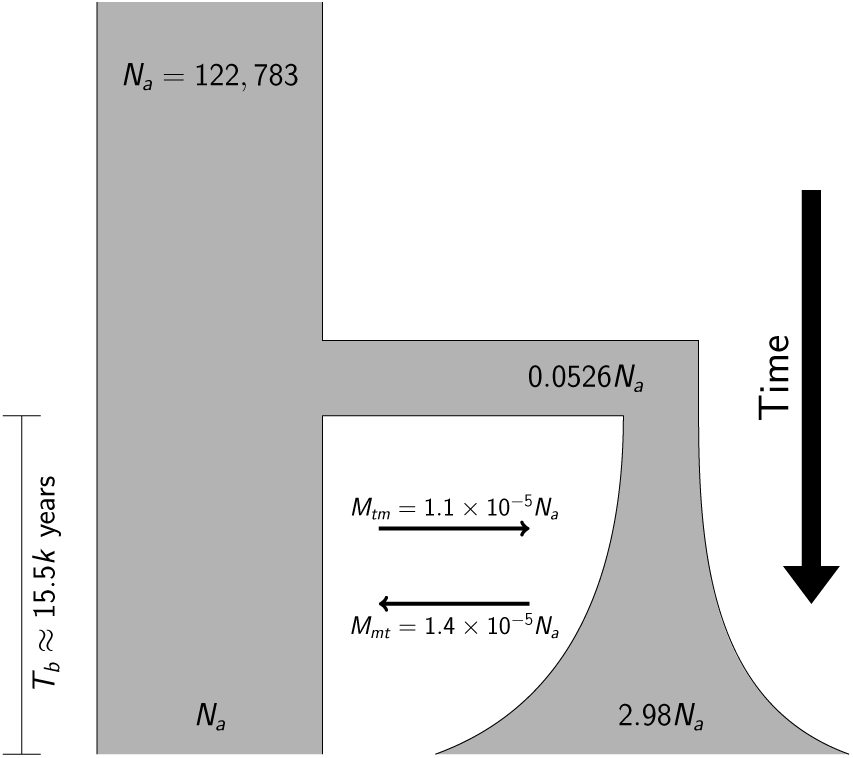
Parameter estimates for a basic bottleneck model of maize domestication. See methods for details.

Because our modest sample size of fully sequenced individuals has limited power to infer recent population expansion, we investigated two alternative approaches for demographic inference. First, we utilized genotyping data from more than 4,000 maize landraces^30^ to estimate the modern maize effective population size. Because rare variants provide the best information about recent effective population sizes^31^, we estimate *N*_*e*_ using a singleton-based estimator^32^ of the population mutation rate *θ* = 4*N*_*e*_*μ* and published values of the mutation rate^33^ (see online methods for details). This yields a much higher estimate of the modern maize effective population size at *N*_*m*_ ≈ 993,000. Finally, we employed a model-free coalescent approach^34^ to estimate population size change using a subset of six genomes each of maize and teosinte. Though this analysis suggests non-equilibrium dynamics for teosinte not included in our initial model, it is nonetheless broadly consistent with the other approaches, identifying population isolation beginning between 10,000 and 15,000 generations ago, a clear domestication bottleneck, and ultimately rapid population expansion in maize to an extremely large extant size of ≈ 10^9^ (Figure S2). Our assessment of the historical demography of maize and teosinte provides context for subsequent analyses of linked selection.

### Hard sweeps do not explain diversity differences

When selection increases the frequency of a new beneficial mutation, a signature of reduced diversity is left at surrounding linked sites^6^. To evaluate whether patterns of such “hard sweeps” could explain observed differences in diversity between genic and intergenic regions of the genome, we compared diversity around missense and synonymous substitutions between *Tripsacum* and either maize or teosinte. If a substantial proportion of missense mutations have been fixed due to hard sweeps, diversity around these substitutions should be lower than around synonymous substitutions. We observe this pattern around the causative amino acid substitution in the maize domestication locus *tga1* (Figure S1), likely the result of a hard sweep during domestication^35,36^. Genome-wide, however, we observe no differences in diversity at sites near synonymous versus missense substitutions in either maize or teosinte (Figure 3).

**Figure 3:**
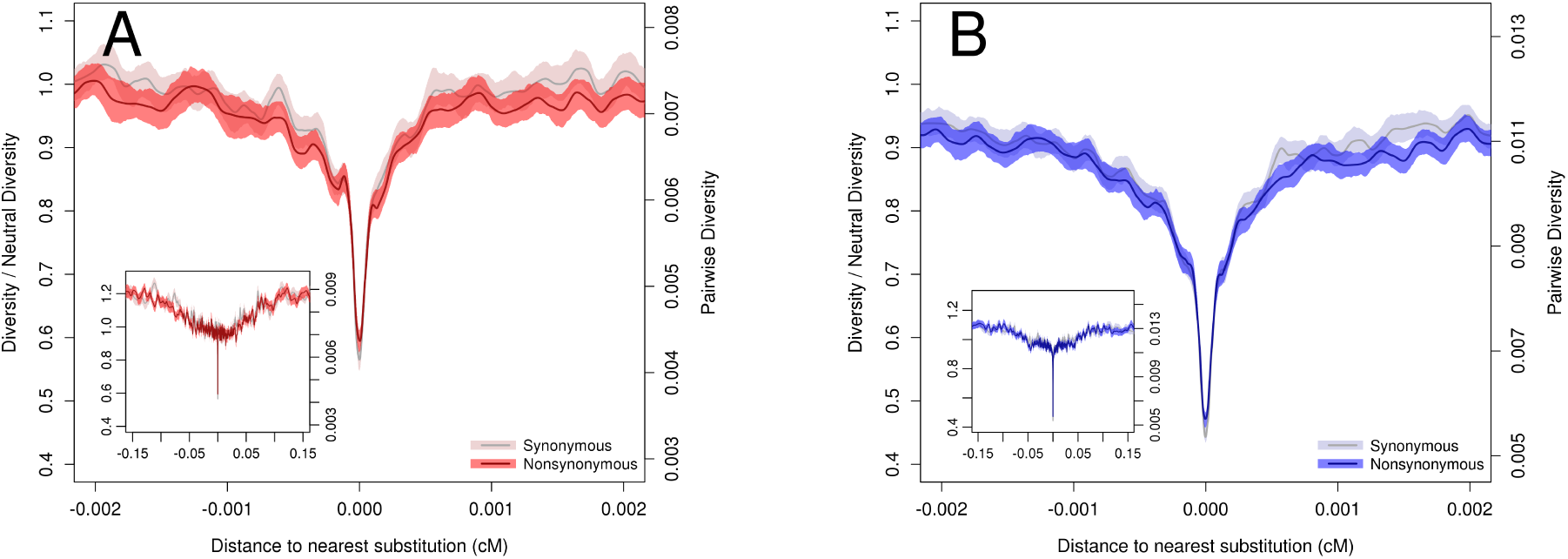
Pairwise diversity surrounding synonymous and missense substitutions in **A** maize and **B** teosinte. Axes show absolute diversity values (right) and values relative to mean nucleotide diversity in windows ≥ 0.01*cM* from a substitution (left). Lines depict a loess curve (span of 0.01) and shading represents bootstrap-based 95% confidence intervals. Inset plots depict a larger range on the x-axis.

Previous analyses have suggested that this approach may have limited power because a relatively high proportion of missense substitutions will be found in genes that, due to weak purifying selection, have higher genetic diversity^37^. To address this concern, we took advantage of genome-wide estimates of evolutionary constraint^38^ calculated using genomic evolutionary rate profile (GERP) scores^39^. We then evaluated substitutions only in subsets of genes in the highest and lowest 10% quantile of mean GERP score, putatively representing genes under the strongest and weakest purifying selection. As expected, we see higher diversity around substitutions in genes under weak purifying selection, but we still find no difference in diversity near synonymous and missense substitutions in either subset of the data (Figure S3). Taken together, these data suggest hard sweeps do not play a major role in patterning genic diversity in either maize or teosinte.

### Diversity is strongly influenced by purifying selection

In the case of purifying or background selection, diversity is reduced in functional regions of the genome via removal of deleterious mutations^9^. We investigated purifying selection in maize and teosinte by evaluating the reduction of diversity around genes. Pairwise diversity is strongly reduced within genes for both maize and teosinte (Figure 4A) but recovers quickly at sites outside of genes, consistent with the low levels of linkage disequilibrium generally observed in these subspecies^26,40^. The reduction in relative diversity is more pronounced in teosinte, reaching lower levels in genes and occurring over a wider region.

**Figure 4:**
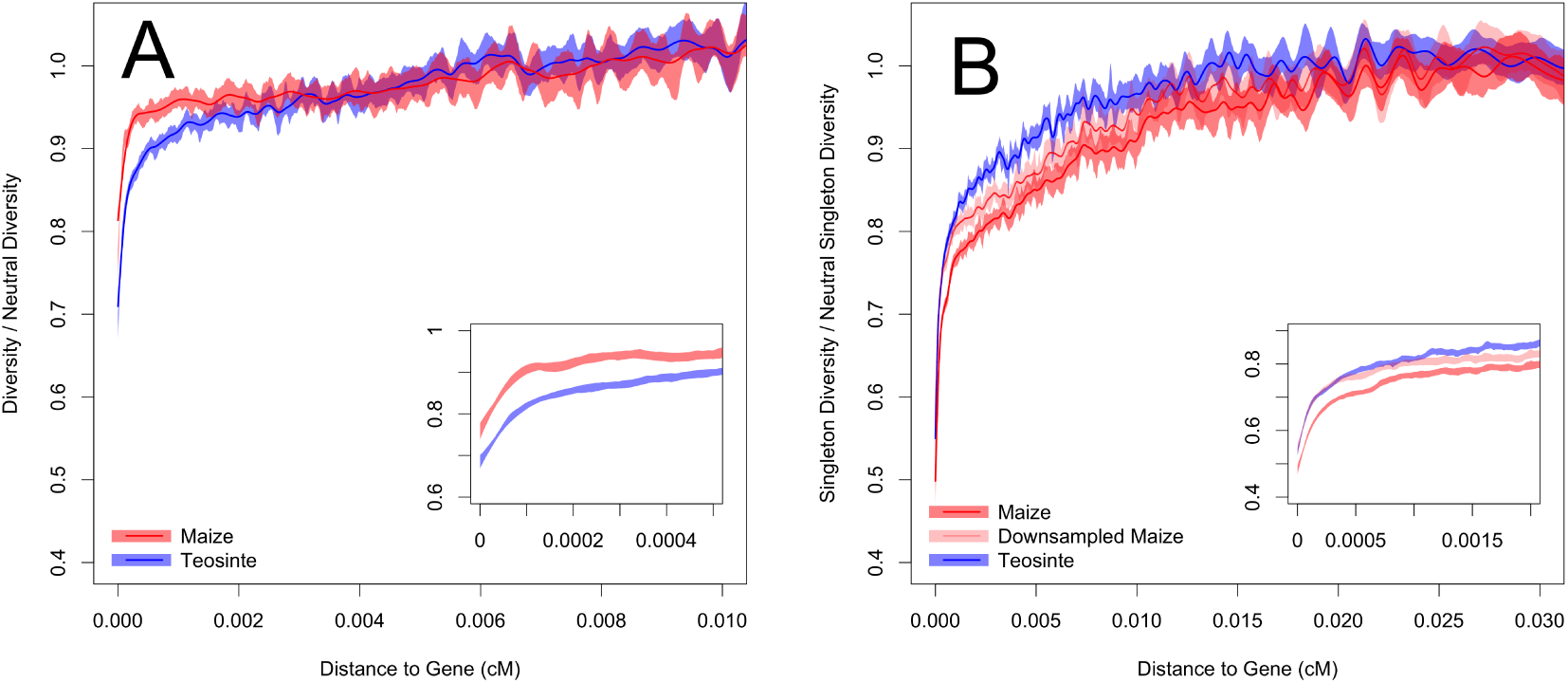
Relative diversity versus distance to nearest gene in maize and teosinte. Shown are **A** pairwise nucleotide diversity and **B** singleton diversity. Relative diversity is calculated compared to the mean diversity in windows ≥ 0.01*cM* or ≥ 0.02*cM* from the nearest gene for pairwise diversity and singletons, respectively. Lines depict cubic smoothing splines with smoothing parameters chosen via generalized cross validation and shading depicts bootstrap-based 95% confidence intervals. Inset plots depict a smaller range on the x-axis.

Our previous comparison of synonymous and missense substitutions has low power to detect the effects of selection acting on multiple beneficial mutations or standing genetic variation, because in such cases diversity around the substitution may be reduced to a lesser degree^41,42^. Nonetheless, such “soft sweeps” are still expected to occur more frequently in functional regions of the genome and could provide an alternative explanation to purifying selection for the observed reduction of diversity at linked sites in genes. To test this possibility, we performed a genome-wide scan for selection using the H12 statistic, a method expected to be sensitive to both hard and soft sweeps^43^. Qualitative differences between maize and teosinte in patterns of diversity within and outside of genes remained unchanged even after removing genes in the top 20% quantile of H12 (Figure S7A). We interpret these combined results as suggesting that purifying selection has predominantly shaped diversity near genes and left a more pronounced signature in the teosinte genome due to the increased efficacy of selection resulting from differences in long-term effective population size.

### Population expansion leads to stronger purifying selection in modern maize

Motivated by the rapid post-domestication expansion of maize evident in our demographic analyses, we reasoned that low-frequency — and thus younger — polymorphisms might show patterns distinct from pairwise diversity, which is determined primarily by intermediate frequency — therefore comparably older — alleles. Singleton diversity around missense and synonymous substitutions (Figure S4) appears nearly identical to results from pairwise diversity (Figure 3), providing little support for a substantial recent increase in the number or strength of hard sweeps occurring in maize.

In contrast, we observe a significant shift in the effects of purifying selection: singleton polymorphisms are more strongly reduced in and near genes in maize than in teosinte, even after downsampling our maize data to account for differences in sample size (Figure 4B). This result is the opposite of the pattern observed for π, where teosinte demonstrated a stronger reduction of diversity in and around genes than did maize. As before, this relationship remained after we removed the 20% of genes with the highest H12 values (Figure S7). While direct comparison of pairwise and singleton diversity within taxa is consistent with non-equilibrium dynamics in teosinte, these too reveal much stronger differences in maize (Figure S5) and mirror results from simulations of purifying selection (Figure S6).

## DISCUSSION

### Demography of domestication

Although a number of authors have investigated the demography of maize domestication^23,24,25^, these efforts relied on data only from genic regions of the genome and made a number of limiting assumptions about the demographic model. We show that diversity within genes has been strongly reduced by the effects of linked selection, such that even synonymous polymorphisms in genes are not representative of diversity at unconstrained sites. This implies that genic polymorphism data are unable to tell the complete or accurate demographic history of maize, but the rapid recovery of diversity outside of genes demonstrates that sites far from genes can be reasonably used for demographic inference. Furthermore, by utilizing the full joint SFS, we are able to estimate population growth, gene flow, and the strength of the domestication bottleneck without making assumptions about its duration. This model paves the way for future work on the demography of domestication, evaluating for example the significance of differences in gene flow estimated here or removing assumptions about demographic history in teosinte.

One surprising result from our model is the estimated divergence time of maize and teosinte approximately 15, 000 generations before present. While this appears to conflict with archaeological estimates^44^, we emphasize that this estimate reflects the fact that the genetic split between populations likely preceded anatomical changes that can be identified in the archaeological record. We also note that our result may be inflated due to population structure, as our geographically diverse sample of teosinte may include populations diverged from those that gave rise to maize.

The estimated bottleneck of ≈ 5% of the ancestral teosinte population seems low given that maize landraces exhibit ≈ 80% of the diversity of teosinte^28^, but our model suggests that the effects of the bottleneck on diversity are likely ameliorated by both gene flow and rapid population growth (Figure 2). Although we estimate that the modern effective size of maize is larger than teosinte, the small size of our sample reduces our power to identify the low frequency alleles most sensitive to rapid population growth^31, and our model is unable to incorporate growth faster than exponential. Both alternative approaches we employ estimate a much larger modern effective size of maize in the range of ≈ 10^6^ – 10^9^, an order of magnitude or more than the current size of teosinte. Census data suggest these estimates are plausible: there are 47.9 million ha of open-pollinated maize in production^45^, likely planted at a density of ≈ 25, 000 individuals per hectare46^. Assuming the effective size is only ≈ 0.4% of the census size (i.e. 1 ear for every 1000 male plants), this still implies a modern effective population size of more than four billion. While these genetic and census estimates are likely inaccurate, all of the evidence points to the fact that the modern effective size of maize is extremely large.

### Hard sweeps do not shape genome-wide diversity in maize

Our findings demonstrate that classic hard selective sweeps have not contributed substantially to genome-wide patterns of diversity in maize, a result we show is robust to concerns about power due to the effects of purifying selection^37^. Although our approach ignores the potential for hard sweeps in noncoding regions of the genome, a growing body of evidence argues against hard sweeps as the prevalent mode of selection shaping maize variability. Among well-characterized domestication loci, only the gene *tga1* shows evidence of a hard sweep on a missense mutation^36^, while published data for several loci are consistent with soft sweeps from standing variation^47,48^ or multiple mutations^49^. Moreover, genome-wide studies of domestication^28^, local adaptation^50^ and modern breeding^51,52^ all support the importance of standing variation as primary sources of adaptive variation. Soft sweeps are expected to be common when 2*N*_*e*_*μ*_*b*_ ≥ 1, where *μ*_*b*_ is the mutation rate of beneficial alleles with selection coefficient *s*_*b*_^42^. Assuming a mutation rate of 3 × 10^−8^^33^ and that on the order of ≈ 1 – 5% of mutations are beneficial^53^, this implies that soft sweeps should be common in both maize and teosinte for mutational targets >> 10*kb* — a plausible size for quantitative traits or for regulatory evolution targeting genes with large up‐ or down-stream control regions^47^ e.g., Indeed, many adaptive traits in both maize^54^ and teosinte^55^ are highly quantitative, and adaptation in both maize^28^ and teosinte^56^ has involved selection on regulatory variation.

The absence of evidence for a genome-wide impact of hard sweeps in coding regions differs markedly from observations in *Drosophila*^57^ and *Capsella*^58^, but is consistent with data from humans^59,60^. Comparisons of the estimated percentages of nonsynonmyous substitutions fixed by natural selection^10,58,61,62^ give similar results. While differences in long-term *N*_*e*_ likely explains some of the observed variation across species, we see little change in the importance of hard sweeps in genes in singleton diversity in modern maize (Figure S4), perhaps suggesting other factors may contribute to these differences as well. One possibility, for example, is that, if mutational target size scales with genome size, the larger genomes of human and maize may offer more opportunities for noncoding loci to contribute to adaptation, with hard sweeps on nonsynonymous variants then playing a relatively smaller role. Support for this idea comes from numerous cases of adaptive transposable element insertion modifying gene regulation in maize^47,63, 64, 65^ and studies of local adaptation that show enrichment for SNPs in regulatory regions in teosinte^56^ and humans^66^ but for nonsynonymous variants in the smaller *Arabidopsis* genome. Our results, for example, are not dissimilar to findings in the comparably-sized mouse genome, where no differences are seen in diversity around nonsynonymous and synonymous substitutions in spite of a large *N*_*e*_ and as many as 80% of adaptive substitutions occuring outside of genes^67^. Future comparative analyses using a common statistical framework (e.g.^14^) and considering additional ecological and life history factors (c.f.^15^) should allow explicit testing of this idea.

### Demography influences the efficiency of purifying selection

One of our more striking findings is that the impact of purifying selection on maize and teosinte qualitatively changed over time. We observe a more pronounced decrease in *π* around genes in teosinte than maize (Figure 4A), but the opposite trend when we evaluate diversity using singleton polymorphisms (Figure 4B). The efficiency of purifying selection is proportional to effective population size^68^, and these results are thus consistent with our demographic analyses which show a domestication bottleneck and smaller long-term *N*_*e*_ in maize^23,24,25,61^ followed by recent rapid expansion and a much larger modern *N*_*e*_. Simple foward-in-time population genetic simulations qualitatively confirm these results, and further suggest that the observed patterns are likely cause by sites under relatively weak purifying selection S6.

Although demographic change affects the efficiency of purifying selection, it may have limited implications for genetic load. Recent population bottlenecks and expansions have increased the relative abundance of rare and deleterious variants in domesticated plants^69,70^ and human populations out of Africa^31,71^, and such variants may play an important role in phenotypic variation^71,72,73^. Nonetheless, demographic history may have little impact on the overall genetic load of populations^74,75^, as decreases in *N*_*e*_ that allow weakly deleterious variants to escape selection also help purge strongly deleterious ones, and the increase of new deleterious mutations in expanding populations is mitigated by their lower initial frequency and the increasing efficiency of purifying selection^75,76,77^.

### Rapid changes in linked selection

Our results demonstrate that consideration of long-term differences in *N*_*e*_ cannot fully capture the dynamic relationship between demography and selection. While a number of authors have tested for selection using methods that explicitly incorporate or are robust to demographic change^62,78,79^ and others have compared estimates of the efficiency of adaptive and purifying selection across species^80^ or populations^81^, previous analyses of the impact of linked selection on genome-wide diversity have relied on single estimates of the effective population size^14,15^. Our results show that demographic change over short periods of time can quickly change the dynamics of linked selection: mutations arising in extant maize populations are much more strongly impacted by the effects of selection on linked sites than would be suggested by analyses using long-term effective population size. As many natural and domesticated populations have undergone considerable demographic change in their recent past, long-term comparisons of *N*_*e*_ are likely not informative about current processes affecting allele frequency trajectories.

## ACKNOWLEDGEMENTS

We are indebted to Graham Coop and Simon Aeschbacher for their constructive input during this study. We thank Robert Bukowski and Qi Sun for providing early-access data from maize HapMap3. Funding was provided by NSF Plant Genome Research Project 1238014 and the USDA-Agricultural Research Service.

## AUTHOR CONTRIBUTIONS

TMB and JRI devised this study. TMB, LW, JRI, and KC analyzed the data. AD performed early-stage simulations. TMB, JRI, and MBH wrote the manuscript.

## ONLINE METHODS

### BASH, R, and Python scripts

All scripts used for analysis are available in an online repository at https://github.com/timbeissinger/Maize-Teo-Scripts.

### Plant materials

We made use of published sequences from inbred accessions of teosinte (*Z. mays* ssp. *parviglumis*) and maize landraces from the Maize HapMap3 panel as part of the Panzea project^26,27,82^. From these data, we removed 4 teosinte individuals that were not ssp. *parviglumis* or appeared as outliers in an initial principal component analysis conducted with the package adegenet^83^ (Figure S8), leaving 13 teosinte and 23 maize that were used for all subsequent analyses (Table S1). We also utilized a single individual of (*Tripsacum dactyloides*) as an outgroup. All bam files are available at /iplant/home/shared/panzea/hapmap3/bam_internal/v3_bams_bwamem.

### Physical and genetic maps

Sequences were mapped to the maize B73 version 3 reference genome^84^ (ftp://ftp.ensemblgenomes.org/pub/plants/release-22/fasta/zea_mays/dna/) as described by^27^. All analyses made use of uniquely mapping reads with mapping quality score ≥ 30 and bases with base quality score ≥ 20; quality scores around indels were adjusted following^85^. We converted physical coordinates to genetic coordinates via linear interpolation of the previously published 1cM resolution NAM genetic map^86^.

### Estimating the site frequency spectrum

We estimated both the genome-wide site frequency spectrum (SFS) as well as a separate SFS for genic (within annotated transcript) and intergenic (≥ 5*kb* from a transcript) regions. We used the biomaRt package^87,88^ of R^89^ to parse annotations from genebuild version 5b of AGPv3. We estimated single population and joint SFS with the software ANGSD^90^, including all positions with at least one aligned read in ≥ 80% of samples in one or both populations. We assumed individuals were fully inbred and treated each line as a single haplotype. Because ANGSD cannot calculate a folded joint SFS, we first polarized SNPs using the maize reference genome and then folded spectra using *δ*a*δ*i^4^.

### Demographic inference

We used the software *δ*a*δ*i^4^ to estimate parameters of a domestication bottleneck from the joint maize-teosinte SFS, using only sites > 5*kb* from a gene to ameliorate the effects of linked selection. To minimize the number of parameters estimated, we employed a simple demographic model which posits a teosinte population of constant effective size *N*_*a*_. At time *T*_*b*_ generations in the past, this population gave rise to a maize population of size *N*_*b*_ which grew exponentially to size *N*_*m*_ in the present (Figure 2). The model includes migration of *M*_*mt*_ individuals each generation from maize to teosinte and *M*_*tm*_ individuals from teosinte to maize. We estimated *N*_*a*_ using *δ*a*δ*i’s estimation of *θ* = 4*N*_*a*_*μ* from the data and a mutation rate of *μ* = 3 × 10^−833^. We estimated all other parameters using 1,000 *δ*a*δ*i optimizations and allowing initial values between runs to be randomly perturbed by a factor of 2. Optimized parameters along with their initial values and upper and lower bounds can be found in table S2. We report parameter estimates from the optimization run with the highest log-likelihood.

We further made use of a large genotyping data set of more than 4,000 partially imputed maize landraces^30^ to estimate the modern maize *N*_*e*_ from singleton counts. We filtered these data to include only SNPs with data in ≥ 1, 500 individuals, and then projected the SFS down to a sample of 500 individuals by sampling each marker without replacement 1,000 times according to the observed allele frequencies. We then estimated *N*_*e*_ from the data assuming *μ* = 3 × 10^−8 33^ and the relation 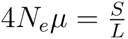^32^, where *S* is the total number of singleton SNPs and *L* is the total number of SNPs in the dataset.

As a final estimate of demography, we employed MSMC^34^ to complement our model-based demographic inference. We used six each of maize and teosinte (BKN022, BKN025, BKN029, BKN030, BKN031, BKN033, TIL01, TIL03, TIL09, TIL10, TIL11 and TIL14), treating each inbred genome as a single haplotype. We called SNPs in ANGSD^90^ using a SNP p-value of 1*e*–6 against a reference genome masked using SNPable (http://lh3lh3.users.sourceforge.net/snpable.shtml). We then removed heterozygous genotypes and filtered sites with a mapping quality < 30, a base quality < 20, or a |*log*_2_(depth)| < 1. We ran MSMC with pattern parameter 20 × 2 + 20 × 4 + 10 × 2 (Figure S2A) for population size inference. To estimate the rate of cross-coalescence we used four maize and four teosinte haplotypes with pattern parameter 20 × 1 + 20 × 2 (Figure S2B).

### Diversity

We made use of the software ANGSD^90^ for diversity calculations and genotype calling. We calculated diversity statistics in maize and teosinte in 1 kb non-overlapping windows using filters as described above for the SFS. We used allele counts to estimate the number of singleton polymorphisms in each window, and used binomial sampling to create a second maize data set down-sampled to have the same number of samples as teosinte. We called genotypes in maize, teosinte, and *Tripsacum* at sites with a SNP p-value < 10^−6^ and when the genotype posterior probability > 0.95. We identified substitutions in maize and teosinte as all sites with a fixed difference with *Tripsacum* and ≤ 20% missing data. Substitutions were classified as synonymous, or missense using the ensembl variant effects predictor^91^. For each window with ≥ 100*bp* of data we computed the genetic distance between the window center and the nearest synonymous and missense substitution as well as the genetic distance to the center of the nearest gene transcript.

### Selection scan

We scanned the genome to identify sites that have experienced recent positive selection using the H12 statistic^43^ in sliding windows of 200 SNPs with a step of 25 SNPs.

### Simulations

We used the program *bneck_selection_ind* included in version 0.4.4 of the forward-in-time population genetic simulation library *fwdpp*^92^ https://github.com/molpopgen/fwdpp]. All simulations used a population mutation rate of *θ* = 20, a population recombination rate of *ρ* = 20, and simulated 150,000 burn-in generations at an ancestral population size of *N*_1_ = 15, 000 to establish equilibrium, after which the population instantly changed to size *N*_2_ and then grew exponentially for 1,000 generations to size *N*_3_. To simulate a constant size population emulating teosinte, we set *N*_2_ = *N*_3_ = 15, 000. For maize we simulated a bottleneck similar to that estimated in Figure 2 by setting *N*_2_ = 750, followed by exponential growth to a large modern population size of *N*_3_ = 150, 000. For each taxon, we performed 1,000 simulations for each of five values of the strength of purifying selection: *s* = (0,10^−6^, 10^−5^, 10^−4^, 10^−3^}. All mutations were assumed to be codominant. To mimic nonsynonymous changes at a coding locus, we assumed that 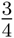 of mutations were selected. We calculated summary statistic across all sites using version 0.3.4 of *msstats* (https://github.com/molpopgen/msstats/releases).

## Supporting Information

**Figure S1:**
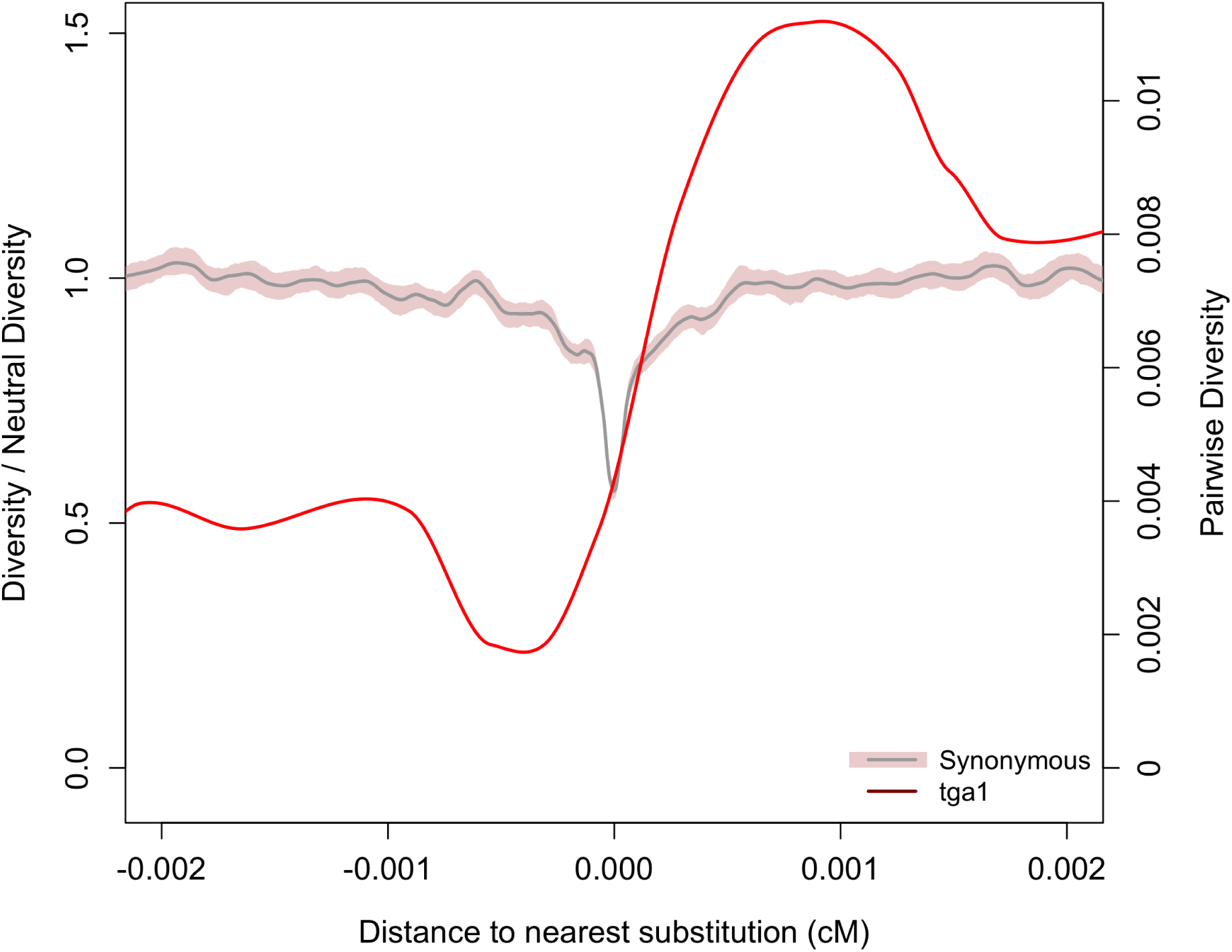
Diversity surrounding the causative substitution at the *tga1* locus.

**Figure S2:**
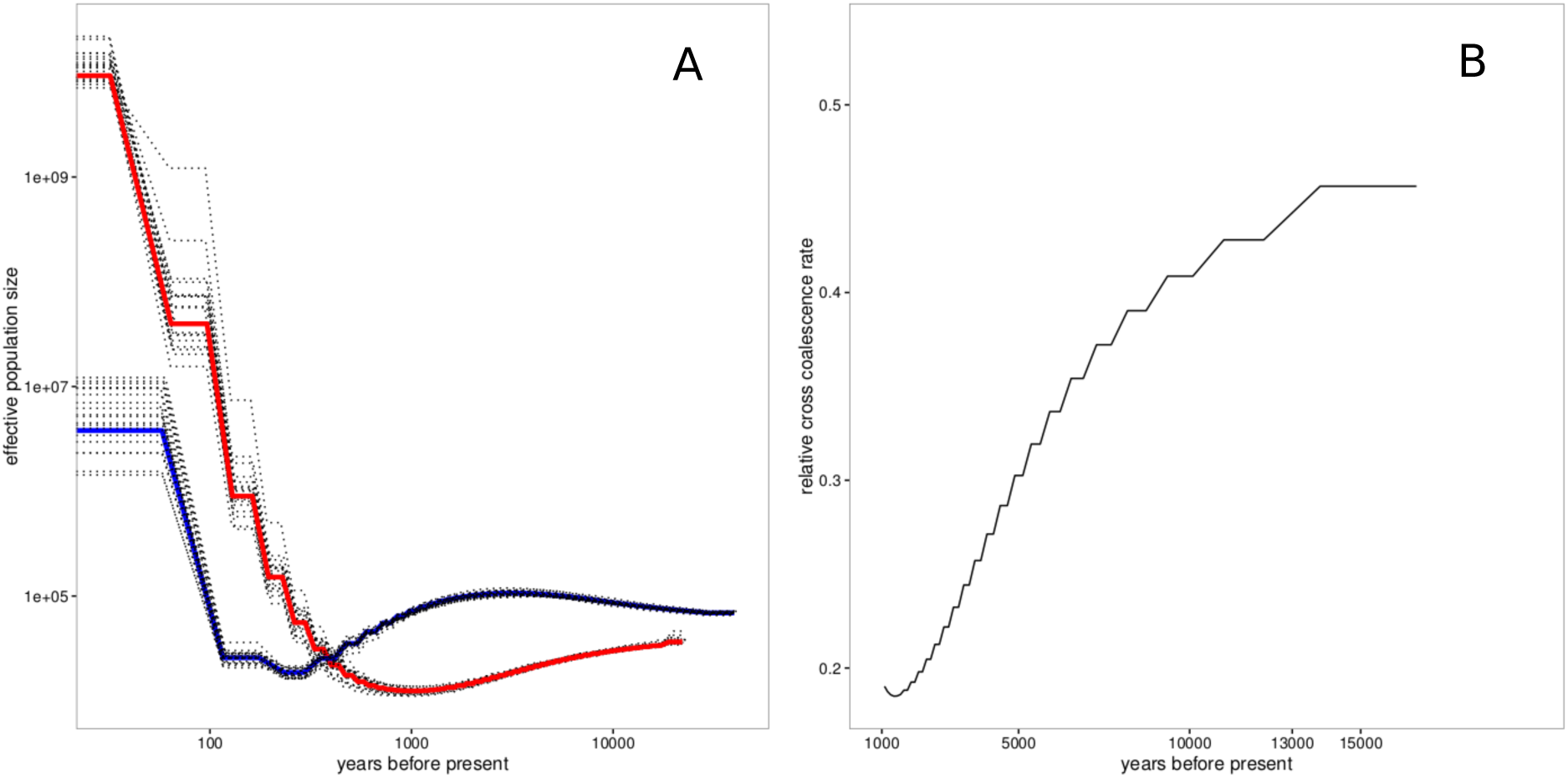
MSMC Analyses. Shown in **A** are effective population size estimates over time. Estimates are depicted as solid lines and boostrap resampling is represented with dotted lines for both maize (red) and teosinte (blue). **B** depticts the relative cross-coalescence rate between maize and teosinte estimated using MSMC. In both panels, time is estimated assuming an annual generation time and a mutation rate of *μ* = 3 × 10^−8^

**Figure S3:**
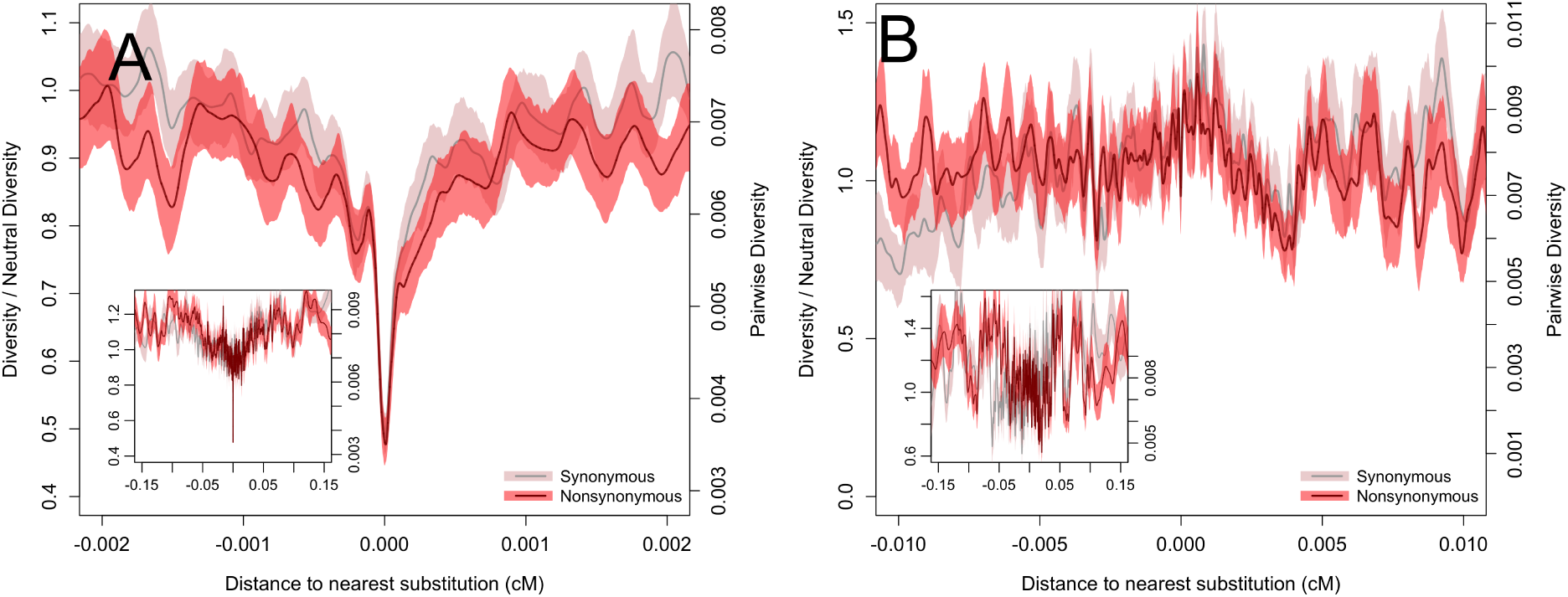
Pairwise diversity surrounding synonymous and nonsynonymous substitutions in maize at **A** highly conserved or **B** unconserved sites. Bootstrap-based 95% confidence intervals are depicted via shading. Inset plots depict a larger range on the x-axis.

**Figure S4:**
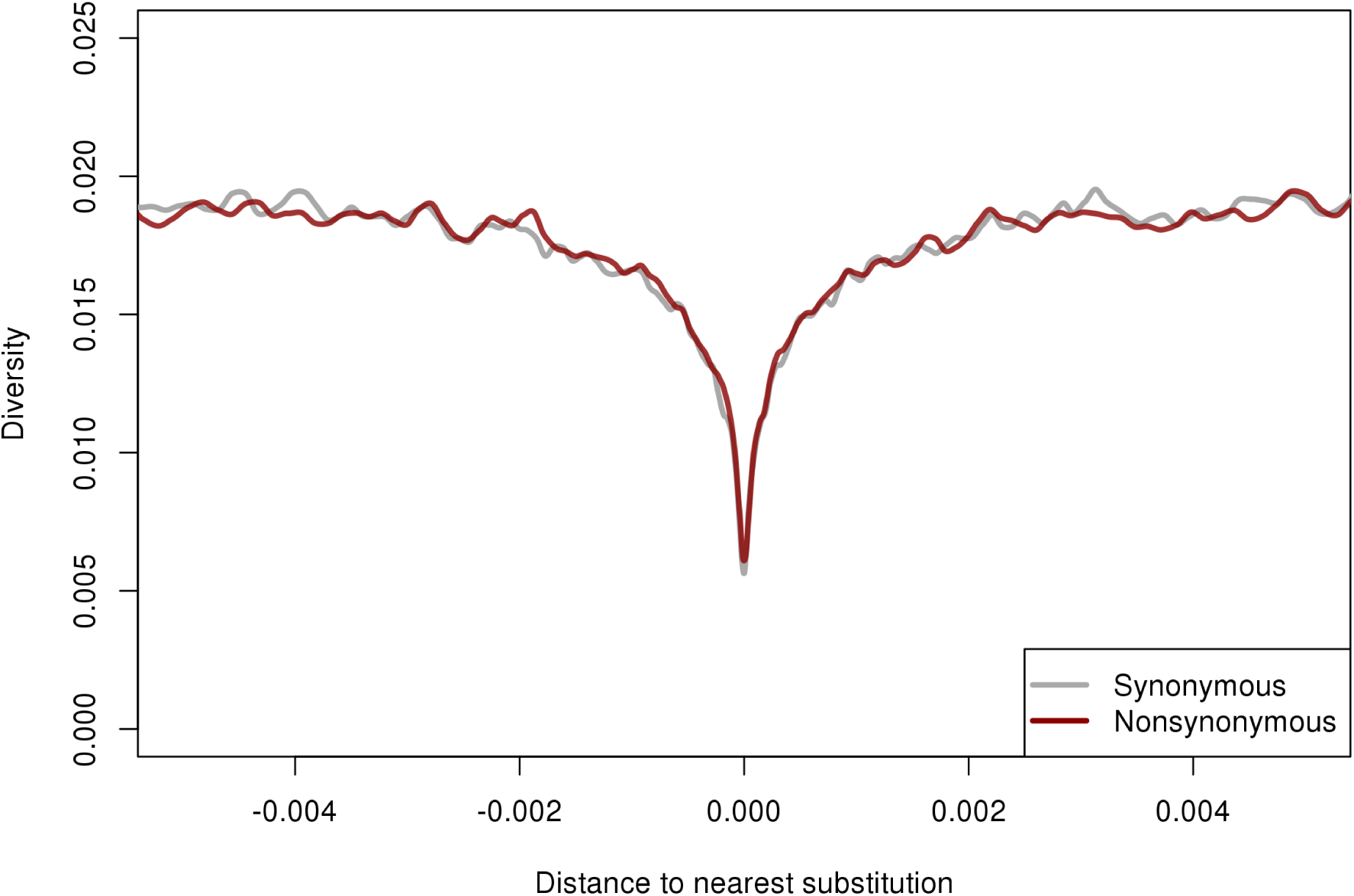
Singleton diversity surrounding synonymous and nonsynonymous substitutions in maize.

**Figure S5:**
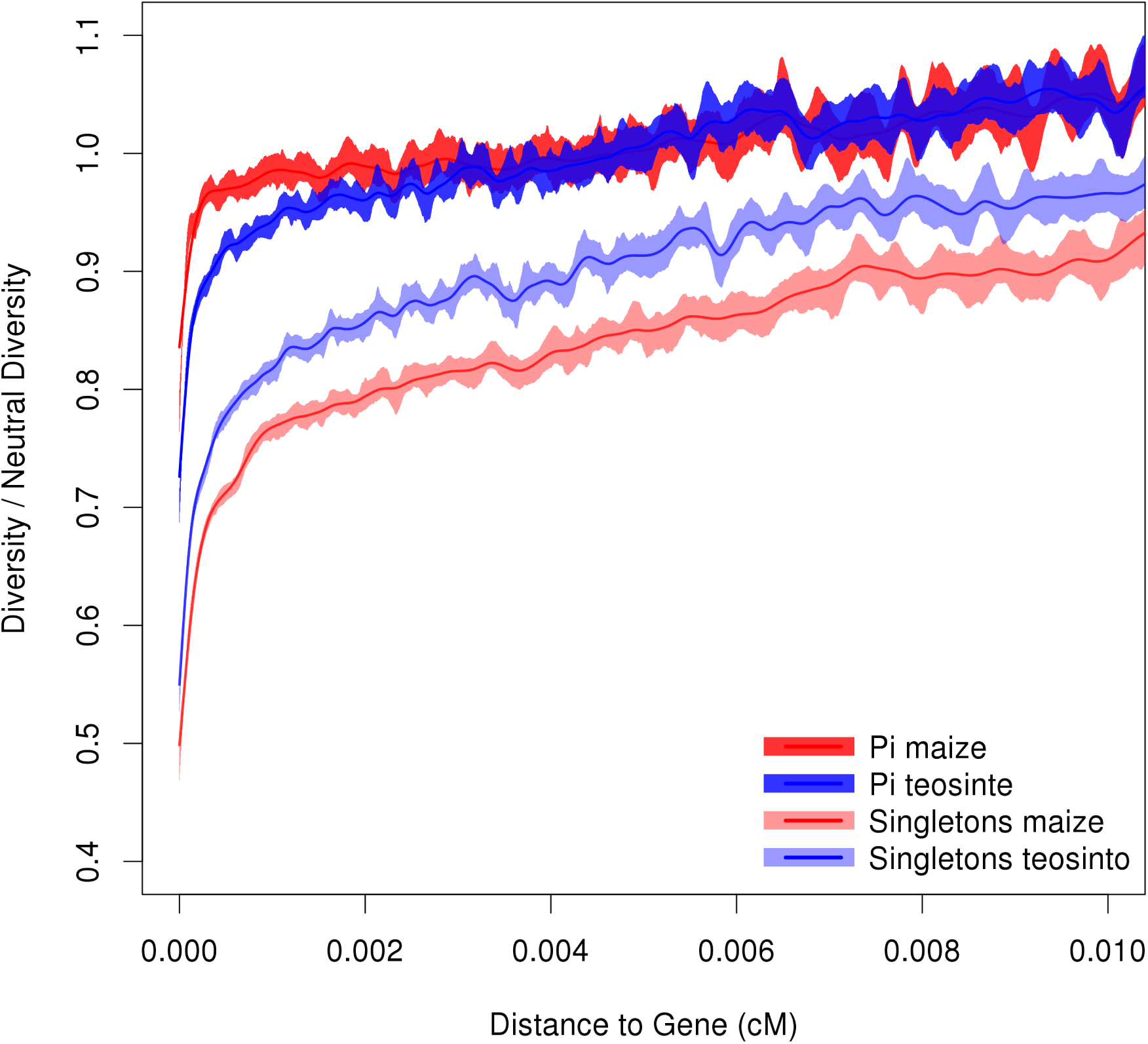
Relative diversity versus distance to nearest gene in maize and teosinte. Relative diversity is calculated by comparing to the mean diversity in all windows ≥ 0.02*cM* from the nearest gene. Lines depict cubic smoothing splines with smoothing parameters chosen via generalized cross validation and shading depicts bootstrap-based 95% confidence intervals.

**Figure S6:**
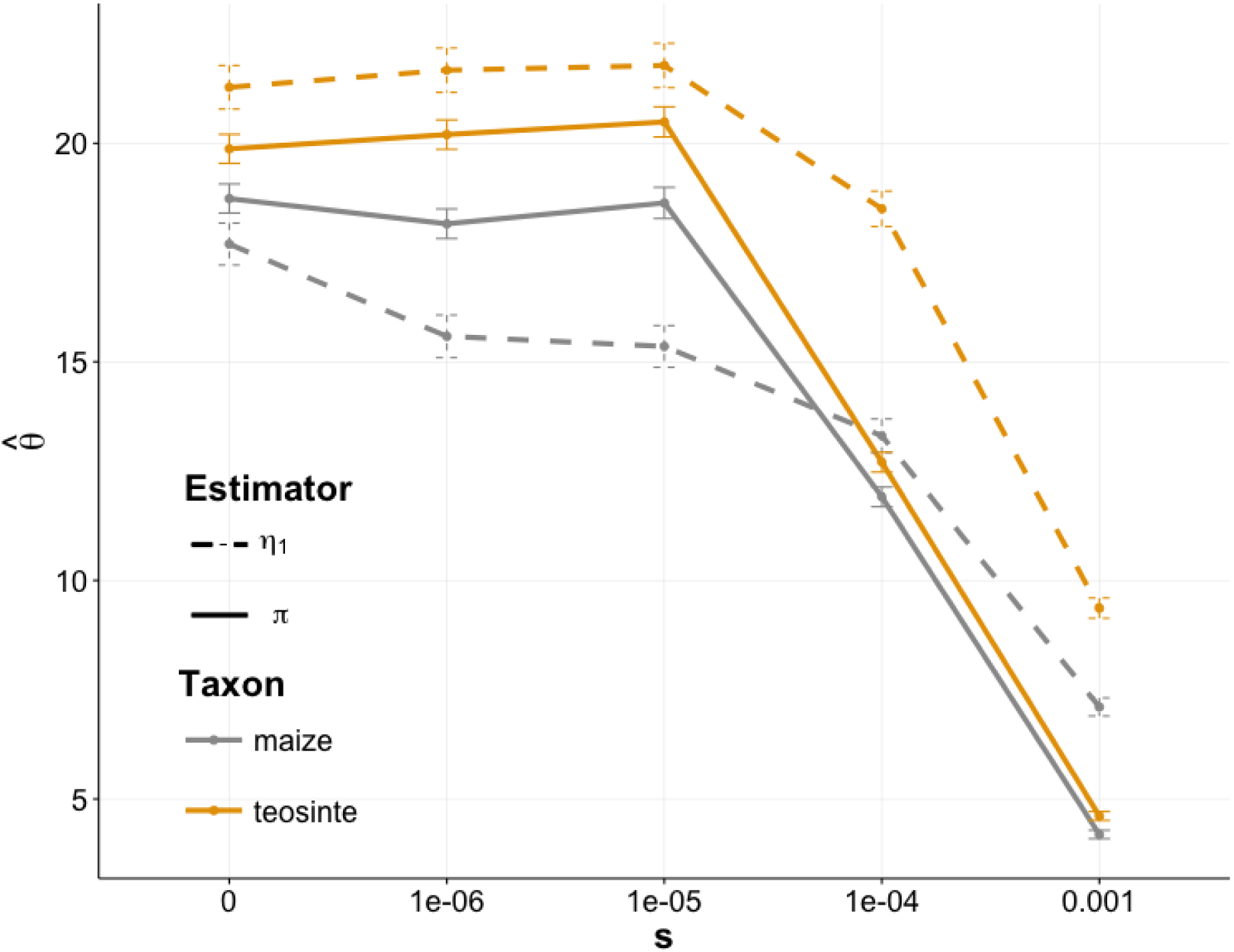
Simulations of diversity statistics in maize and teosinte with varying strengths of purifying selection. Points show the mean (± standard error) of the population mutation rate *θ* estimated by singletons (*η*_1_) and pairwise differences (*π*).

**Figure S7:**
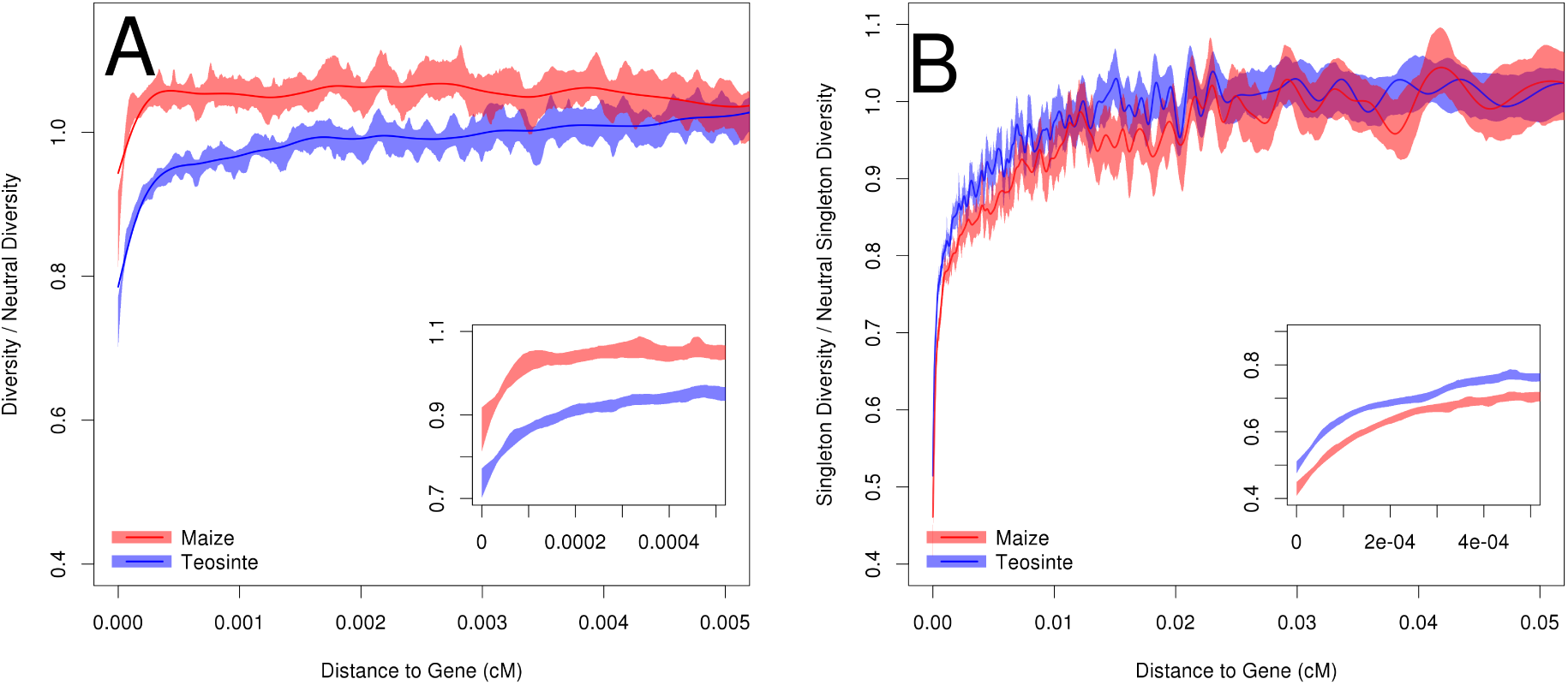
Relative level of diversity versus distance to the nearest gene, in maize and teosinte, based on only sites that do not show evidence of hard or soft sweeps according to H12. Two measures of diversity were investigated. **A** displays pairwise diversity, which is most influenced by intermediate frequency alleles and therefore depicts more ancient evolutionary patterns, and **B** depicts singleton diversity, influenced by rare alleles and thus depicting evolutionary patterns in the recent past. Bootstrap-based 95% confidence intervals are depicted via shading. Inset plots depict a smaller range on the x-axis.

**Figure S8:**
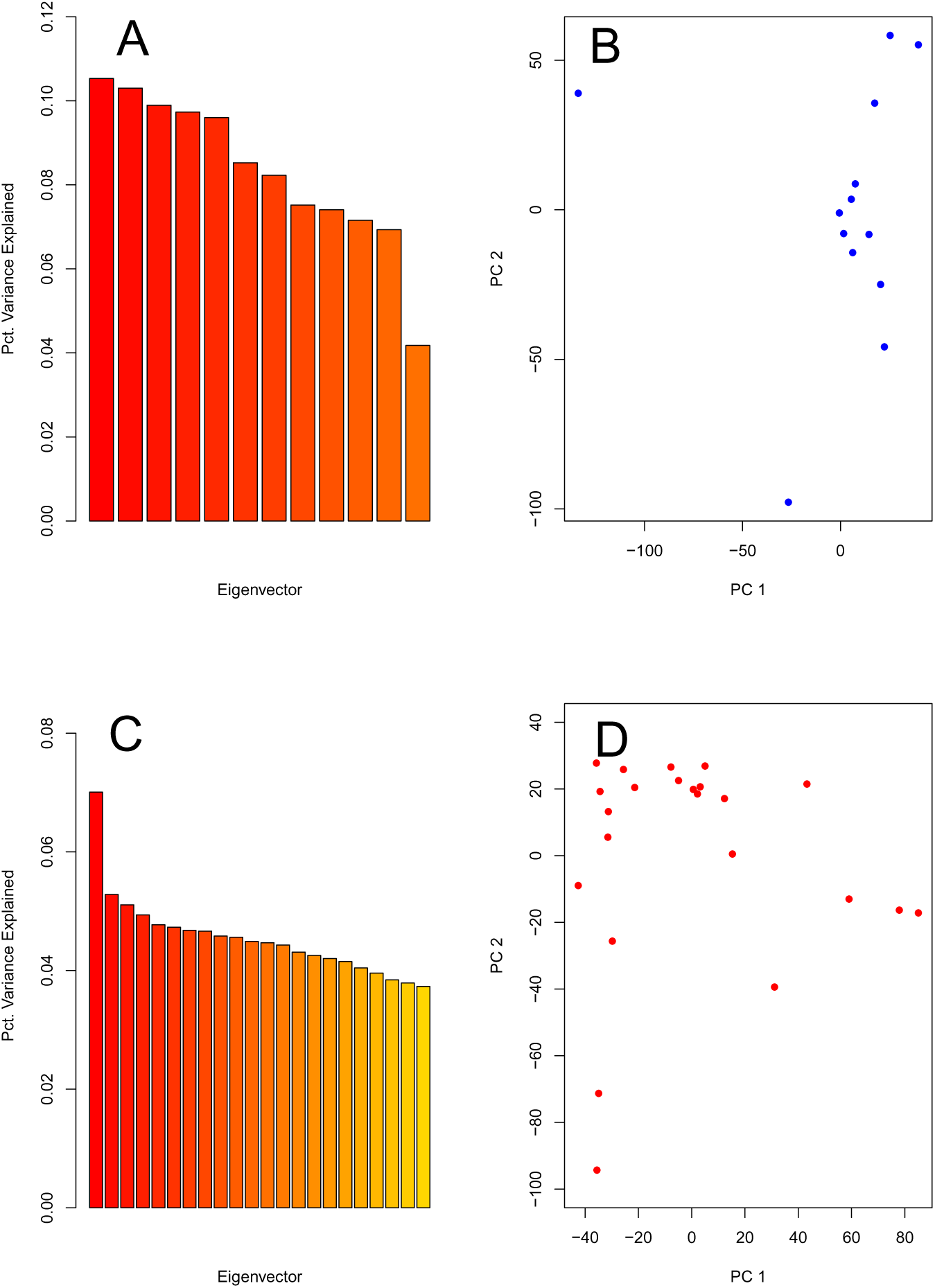
Principal component analysis of teosinte and maize individuals to ensure that no close relatives were inadvertantly included in our study. Plots are based on a random sample of 10,000 SNPs. **A** displays the percentage of total variance explained by each principal component for teosinte, while **B** shows PC1 vs PC2 for all 13 teosinte individuals. Simlarly, **C** depicts the percentage of total variance explained by each principal component for maize, and **D** shows PC1 vs PC2 for all 23 maize individuals.

**Table S1:** A list of maize and teosinte individuals included in this study. Sequencing and details were previously described by^26^

| **Maize** | **Teosinte** |
| --- | --- |
| BKN009 | TIL01 |
| BKN010 | TIL02 |
| BKN011 | TIL03 |
| BKN014 | TIL04-TIP454 |
| BKN015 | TIL07 |
| BKN016 | TIL09 |
| BKN017 | TIL10 |
| BKN018 | TIL11 |
| BKN019 | TIL12 |
| BKN020 | TIL14-TIP498 |
| BKN022 | TIL15 |
| BKN023 | TIL16 |
| BKN025 | TIL17 |
| BKN026 |  |
| BKN027 |  |
| BKN029 |  |
| BKN030 |  |
| BKN031 |  |
| BKN032 |  |
| BKN033 |  |
| BKN034 |  |
| BKN035 |  |
| BKN040 |  |

**Table S2.** Parameters, initial values, and boundaries used for model-fitting with *δαδi*. Parameters are shown in the units utilized by *δαδi*, although in the text simplified units are reported.

| Parameter | Initial value | Upper bound | Lower bound |
| --- | --- | --- | --- |
| $\frac{N_b}{N_a}$ | 0.02 | $1 \times 10^{-7}$ | 2 |
| $\frac{N_m}{N_a}$ | 3 | $1 \times 10^{-7}$ | 200 |
| $\frac{T_b}{2N_a}$ | 0.04 | 0 | 1 |
| $\frac{M_{mt}}{N_a}$ | $1 \times 10^{-10}$ | $1 \times 10^{-7}$ | 0.001 |
| $\frac{M_{tm}}{N_a}$ | $1 \times 10^{-10}$ | $1 \times 10^{-7}$ | 0.001 |

